# Association between Cognitive Function and Large Optic Nerve Cupping, Accounting for Cup-Disc-Ratio Genetic Risk Score

**DOI:** 10.1101/2021.09.30.462651

**Authors:** Sasha Kravets, Rawan Allozi Rupnow, Abhishek Sethi, Mark A. Espeland, Louis R. Pasquale, Stephen R. Rapp, Barbara E. Klein, Stacy M. Meuer, Mary N. Haan, Pauline M. Maki, Joelle A. Hallak, Thasarat Sutabutr Vajaranant

**Affiliations:** Illinois Eye and Ear Infirmary, Department of Ophthalmology and Visual Sciences, University of Illinois at Chicago, Chicago, Illinois, USA; Division of Epidemiology and Biostatistics, School of Public Health, University of Illinois at Chicago, Chicago IL; Department of Medicine, University of Illinois at Chicago, Chicago, IL; Departments of Internal Medicine and Biostatistics and Data Science, Wake Forest University Health Sciences, Winston Salem, North Carolina, USA; Department of Ophthalmology, Icahn School of Medicine at Mount Sinai, New York, New York, USA; Psychiatry and Behavioral Medicine, Wake Forest University Health Sciences, Winston Salem, North Carolina, USA; Department of Ophthalmology and Visual Sciences, University of Wisconsin, Madison, Wisconsin, USA; Department of Epidemiology and Biostatistics, University of California at San Francisco, San Francisco, California, USA; Department of Psychiatry, University of Illinois at Chicago, Chicago, Illinois, USA

## Abstract

**Purpose:** To investigate if accounting for a cup-to-disc ratio (CDR) genetic risk score (GRS) modified the association between large CDR and cognitive function among women.

**Design:** This was a retrospective study using data from the Women’s Health Initiative.

**Methods:** Patients with glaucoma or ocular hypertension were excluded. Large CDR was defined as ≥ 0.6 in either eye. Cognitive function was measured by the Modified Mini-Mental State Examination (3MSE). We used the combined effects from 13 single nucleotide polymorphisms (SNPs) to formulate the GRS for CDR. We used logistic regression to investigate associations between weighted GRS and large CDR, then a linear regression to assess the association between weighted GRS and 3MSE scores, and between weighted GRS, CDR, and 3MSE scores, adjusted for demographic and clinical characteristics.

**Results:** Final analyses included 1,196 White women with mean age of 69.60 ± 3.62 years and 7.27% with large CDR. Mean GRS in women with and without large CDR was 1.51 ± 0.31 vs. 1.41 ± 0.36, respectively (p = 0.004). The odds of large CDR for a one unit increase in GRS was 2.30 (95% CI: (1.22, 4.36), p = 0.011). Adding the CDR GRS in the model with CDR and 3MSE, women with large CDR still had statistically significantly lower 3MSE scores than those without large CDR, yielding a predicted mean difference in 3MSE scores of 0.84 (p = 0.007).

**Conclusions:** Independent of the CDR GRS, women with large CDR had a lower cognitive function.

## Introduction

Glaucoma, currently affecting more than 70 million individuals worldwide, is the leading cause of irreversible blindness.^1^ While its pathogenesis is still poorly understood, older age and elevated intraocular pressure (IOP) are considered to be important risk factors.^2–3^ Glaucoma progression can also occur in patients with normal range IOP, suggesting that other risk factors may be involved in its pathogenesis.^4^ One candidate risk factor is neurodegeneration of the optic nerve and retinal ganglion cells that occurs with normal aging and diseases such as Alzheimer’s disease (AD), which is a leading cause of dementia worldwide.^5^

Several studies have supported a connection between primary open-angle glaucoma (POAG) and AD. Epidemiological studies have shown a higher prevalence of glaucoma among AD patients.^6–7^ Visual difficulties, such as in reading and finding objects, altered depth and color perception, impaired spatial contrast sensitivity or difficulties in perceiving structure from motion have also been noted in AD patients.^8^ In addition, Optical Coherence Tomography (OCT) studies in AD patients show preferential retinal nerve fiber layer (RNFL) thinning in the superior quadrant of the optic nerve which has a preponderance of large axons, resembling a pattern seen in glaucomatous eyes.^9–10^ Degeneration of the optic nerve leads to cupping, which can be quantified by cup-to-disc ratio (CDR) and which serves as a biomarker for glaucomatous neuropathy. We previously reported that a large CDR (i.e., greater or equal to 0.6) was associated with lower cognitive function, in White post-menopausal women aged 65 years and older without glaucoma or ocular hypertension.^11^

Previous Genome Wide Association Studies (GWAS), including a genome-wide meta-analysis, have identified multiple loci associated with vertical CDR.^12^ While multiple loci have been identified, individually each loci provides limited information. For this reason, a genetic risk score (GRS), combining information from multiple genetic variants into a single measure, has been used to study the genetic risk factors for POAG as well as vertical CDR. Furthermore, previous studies have shown that genetic variants can vary across racial groups. To account for these variations, in our study, we focused on White patients, specifically elderly White women aged 65 years and older.

In this study, our goals were to (1) develop a GRS for CDR and (2) determine if the GRS modified the previously identified relationship between large CDR and decreased cognitive function in cognitively intact women aged 65 years and older presenting without glaucoma or ocular hypertension. This work provides insight into the possible connections between genetics and neurodegenerative changes of the optic nerve and brain.

## Materials & Methods

The Institutional Review Board at the University of Illinois at Chicago waived IRB approval for this secondary data analysis of de-identified data from the WHI. This study adhered to the Declaration of Helsinki and all federal and state laws.

### Data Source

The Women’s Health Initiative (WHI) is a series of clinical trials and observational studies funded by the National Heart, Lung, and Blood Institute from 1991 planned through 2027. The original WHI study, starting in 1991, enrolled over 68,000 postmenopausal women between the ages of 50 and 79 and was split into three separate double blinded randomized control trials. The trials investigated the effects of hormone therapy, dietary modification, and calcium and vitamin D supplements for prevention of heart disease, osteoporosis and its related effects, and associated risks for breast and colorectal cancer. Participants in the trials were stratified by hysterectomy status: with a uterus versus without a uterus. Those with a uterus were randomized to either placebo or estrogen plus progestin, and those without a uterus were randomized to either placebo or estrogen-alone. The WHI trial is registered at https://www.clinicaltrials.gov (NCT00000611).

Within the WHI Hormone Therapy trials, we obtained data from two ancillary studies: the WHI-Sight Exam (WHISE), conducted from 2000 to 2002, and the WHI Memory Study (WHIMS), conducted between 1996 and 2007. The WHISE study recruited participants approximately five years after the randomization into the WHI hormone trial. The goal of this study was to investigate the effect of hormone therapy on age-related macular degeneration. Overall, 4,347 participants consented and completed enrollment with fundus photographs of at least one eye available. To determine CDR, the outer edges of the vertical margin of the optic disc and cup were compared with a series of cup-to-disc standards of increasing size. If the ratio fell between 2 standards the smaller ratio was chosen. Within the WHISE study, vertical CDR was classified as a binary variable, where large CDR was defined as CDR ≥ 0.6 in at least one eye. The WHIMS study investigated the relationship between the effects of hormone therapy and the development and progression of memory loss in women. Participants underwent a baseline global cognitive function test that was repeated annually for ten years. Global cognitive function was assessed by the Modified Mini-Mental State Examination (3MSE) scores. 3MSE scores ranged from 0 to 100, where higher scores indicated better cognitive function.

Within the WHI, genetic data was available from the WHI harmonized and imputed data, which combined genetic data from six genome wide association studies (GWASs) within the WHI Clinical Trials and Observational Studies. The harmonization of over 30,000 samples involved alignment to the same reference panel, imputation to 1000 genomes, identity by descent (IBD) analysis for identification of genetically related individuals, and principal component analysis (PCA) for comparison with self-reported ethnicity. Genotypic quality control (QC) was performed by study and included evaluation of minimal sample call rate, minimal SNP call rate, Hardy-Weinberg equilibrium, and minor allele frequencies. Details regarding QC, harmonization, and imputation of the genetic data can be found from dbGaP at http://www.ncbi.nlm.nih.gov/sites/entrez?db=gap through dbGaP accession phs000746.v2.p3.

### Sample Selection

Participants were selected for analysis if they were included in the WHISE or WHIMS studies, and had genetic data within the harmonized imputation dataset. The harmonized imputation dataset contains 3,111 genetic samples for 2,708 participants of the WHISE study. The majority, 2,339 out of the 2,708 (86%) self-identified as White. Additionally, 12 Black or African American participants, 0 Hispanic/Latino, and 5 other race participants had large CDR. While there were cognitive differences between White vs non-White (mean ± SD 3MSE score of 96.94 ± 3.33 vs 93.17 ± 6.43, p-value<0.001, respectively), given the small distribution of CDR among non-White participants, and that SNPs were only available for participants of European ancestry, the study was limited to White participants. For White subjects with duplicate samples, we included the genetic sample from the GWAS platform with a larger number of SNPs available. We identified three pairs of first-degree relatives and removed the subjects with genetic data from the smaller GWAS platform. Furthermore, 28 subjects had missing PCA data or CDR status. We did not identify inconsistent ethnicity from self-reported ethnicity among subjects with PCA data. After merging the 2,336 patients with genetic information with data from WHISE and WHIMS, 1,196 independent WHISE subjects from the WHI Harmonized Imputation dataset were included in the final analysis. The study schema is depicted in Figure 1.

**Fig 1.**
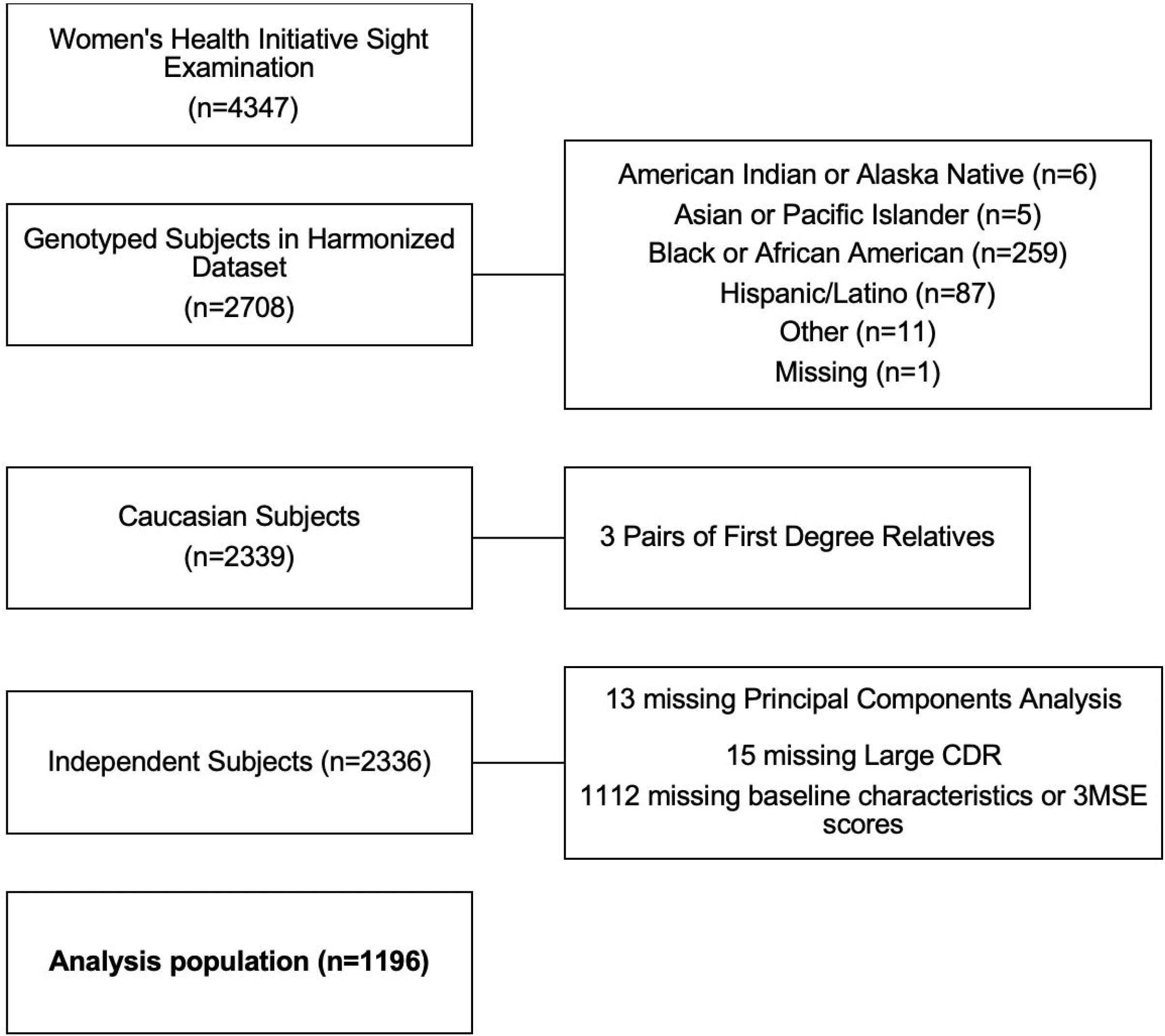
Schema: diagram of analysis of patient population.

### Cup-Disc Ratio Genetic Risk Score

We selected 18 candidate single nucleotide polymorphisms (SNPs) associated with vertical CDR in White patients by a meta-analysis reported in Springelkamp et al.^12^ Imputation quality was assessed by R^2^, where R^2^ is the estimated value of the squared correlation between imputed genotypes and true, unobserved genotypes. The imputation quality for rs1345 (chromosome 11, build 37 position 65337251) in one study was low (R^2^=0.56). The remaining 17 out of the 18 SNPs had high imputation quality (R^2^ > 0.89) across all studies (Table 1). For the 17 high imputation quality SNPs, we performed a logistic regression between large CDR and each SNP under the assumption of an additive model for the effect of the risk allele adjusted for age and the first two principal components (PC-1 and PC-2). There was concordance on the risk allele for 13 out of the 17 SNPs as reported by Springelkamp et al. These 13 SNPs were used to formulate the CDR GRS (Table 2).

**Table 1.**
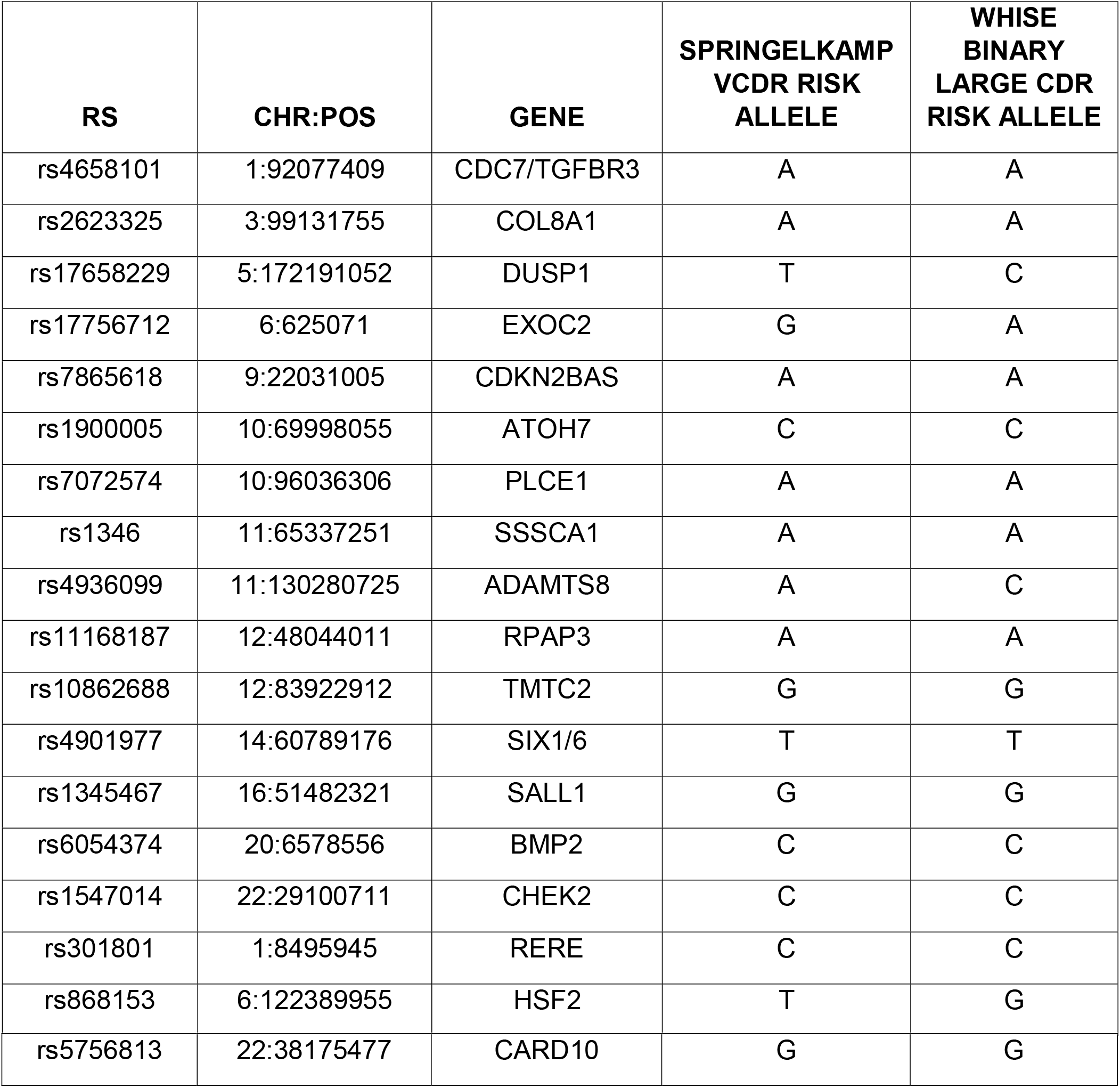
18 candidate SNPs associated with vertical cup-to-disc ratio.

**Table 2.** Final 13 SNPs included in Weighted Genetic Risk Score based on high imputation quality (R^2^) and matching risk alleles.

| SNP | Gene | Weight |
| --- | --- | --- |
| 1:92077409 | CDC7/TGFBR3 | 0.0002 |
| 3:99131755 | COL8A1 | 0.0337 |
| 9:22031005 | CDKN2BAS | 0.3245 |
| 10:69998055 | ATOH7 | 0.0852 |
| 10:96036306 | PLCE1 | 0.0207 |
| 12:48044011 | RPAP3 | 0.0523 |
| 12:83922912 | TMTC2 | 0.1488 |
| 14:60789176 | SIX1/6 | 0.1462 |
| 16:51482321 | SALL1 | 0.0544 |
| 20:6578556 | BMP2 | 0.2726 |
| 22:29100711 | CHEK2 | 0.0702 |
| 1:8495945 | RERE | 0.0966 |
| 22:38175477 | CARD10 | 0.0537 |

We used the log odds ratio (OR) estimated from the single SNP logistic regression model adjusted for age, PC-1, and PC-2 to form the GRS. For *m* patients, we define *G_i_* as the number of risk alleles for the *i*th SNP, and log *OR_i_* as the log OR for the *i*th SNP from the single SNP logistic regression model. The GRS is defined as 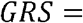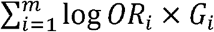. A weighted GRS was constructed by multiplying the vertical CDR-increasing allele by the effect size previously reported.

### Statistical Analysis

Demographic and baseline clinical variables were obtained from this data set including, education, smoking, diabetes, body mass index (BMI), cardiovascular disease, diabetic retinopathy, and hormone therapy randomization. T-tests or Wilcoxon rank sum tests were used to test associations for continuous variables, and chi-square tests or Fisher’s exact test were used to test associations with categorical variables.

We used logistic regression to evaluate the cross-sectional association of the weighted GRS with large CDR. We used linear regression models to evaluate the association between weighted GRS and 3MSE scores, and between weighted GRS, CDR, and 3MSE, both models adjusted for age, education, smoking, diabetes, body mass index, cardiovascular disease, diabetic retinopathy, and hormone therapy randomization. Specifically, we investigated models using the previously mentioned covariates, including an interaction between GRS and CDR, with 3MSE scores as the outcome of interest. As 3MSE scores were not normally distributed, we used a log-transformed function of scores, log (102-3MSE). All analyses were conducted using R (R Core Team (2019). R: URL https://www.R-project.org/).

## Results

Final analyses included 1,196 White women with baseline CDR, demographic and clinical characteristic data, 3MSE scores, and weighted GRS. The mean age (± SD) was 69.60 ± 3.62 years. 87 of 1,196 (7.27%) women had large CDR. Table 3 shows the distribution of the baseline demographic and clinical characteristic variables. Figure 2 shows the distribution of GRS by large CDR status. The mean GRS in women with and without a large CDR was 1.51 ± 0.31 vs 1.41 ± 0.36 (p-value=0.004).

**Table 3.** Patient Demographic and Clinical Characteristics.

| **Variable** | **Summary Statistics** |
| --- | --- |
| **Number of Patients, n** | 1196 |
| **Age at Enrollment, Mean (SD) [Min, Max]** | 69.6 (3.62) [63,79] |
| **Education, n(%)** |  |
| No High School | 50 (4.18) |
| High School | 294 (24.58) |
| Post High School | 463 (38.71) |
| College Graduate | 389 (32.53) |
| **Cigarette smoking, n(%)** |  |
| Never Smoked | 690 (57.69) |
| Former Smoker | 440 (36.79) |
| Current Smoker | 66 (5.52) |
| **BMI, n(%)** |  |
| Underweight | 9 (0.75) |
| Normal | 348 (29.1) |
| Overweight | 412 (34.45) |
| Obesity Class 1 | 288 (24.08) |
| Obesity Class 2 | 98 (8.19) |
| Extreme Obesity | 41 (3.43) |
| **Diabetes, n(%)** |  |
| No | 1101 (92.06) |
| Yes | 95 (7.94) |
| **Prior Cardiovascular Disease, n(%)** |  |
| No | 1009 (84.36) |
| Yes | 187 (15.64) |
| **Diabetic Retinopathy, n(%)** |  |
| No | 1168 (97.66) |
| Yes | 28 (2.34) |

| **Hormone therapy assignment, n(%)** |  |
| --- | --- |
| E-alone intervention | 213 (17.81) |
| E-alone control | 240 (20.07) |
| E+P intervention | 369 (30.85) |
| E+P control | 374 (31.27) |
| **Large Cup-to-Disc Ratio, n(%)** |  |
| Without | 1109 (92.73) |
| With | 87 (7.27) |

**Fig 2.**
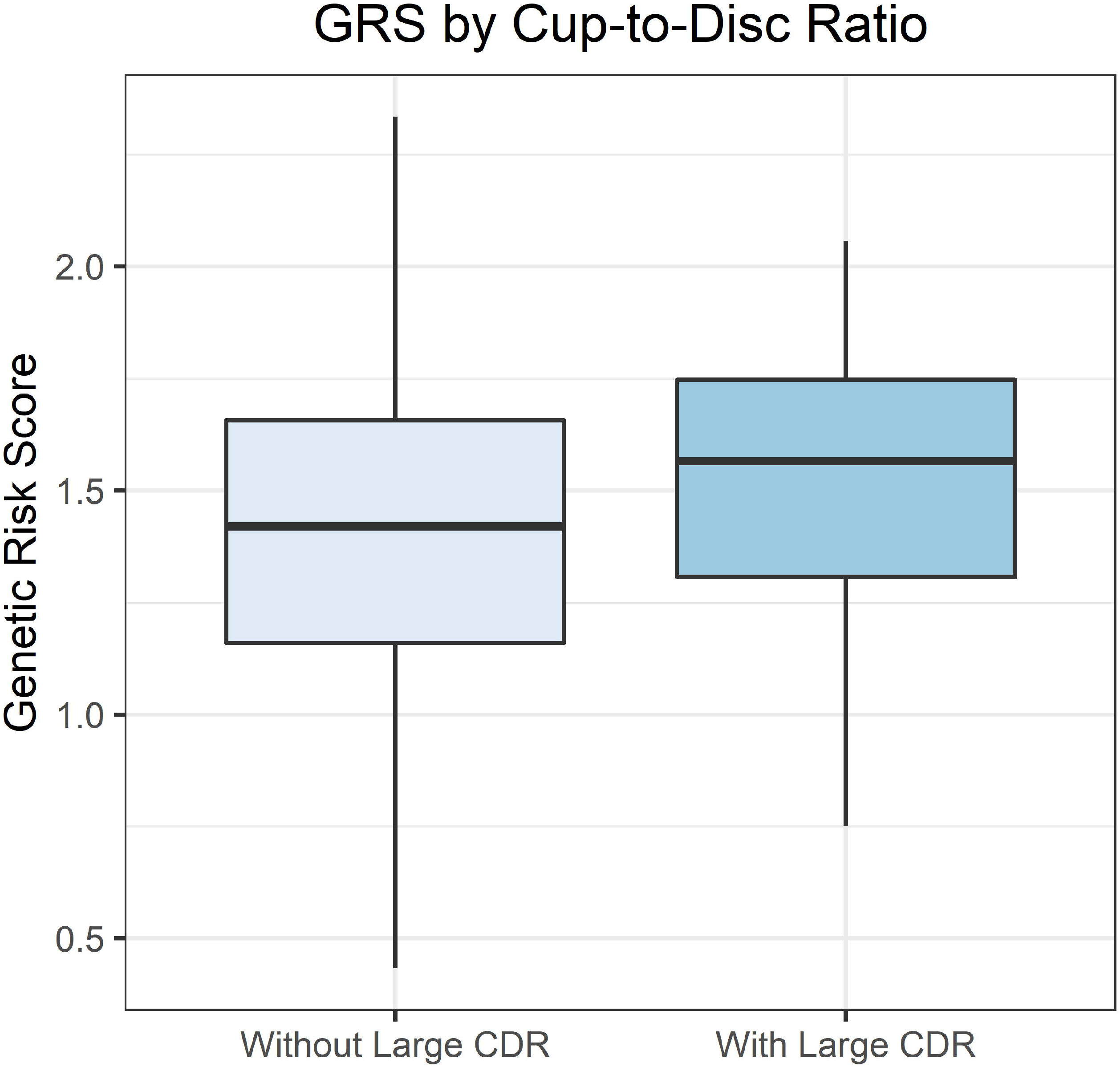
Comparison of Weighted Genetic Risk Score by Large Optic Nerve Cupping.

In the model adjusted for age, education, smoking, diabetes, BMI, cardiovascular disease, diabetic retinopathy, and hormone therapy randomization, White women with large CDR had statistically significantly lower 3MSE scores, compared to White women without large CDR, yielding a predicted mean difference in 3MSE scores of 0.83 (p-value = 0.007).

In a logistic model adjusted for age, PC-1, and PC-2, we found that weighted GRS was associated with large CDR. The odds of large CDR for a one-unit greater GRS was 2.30 (95% CI: (1.21, 4.35), p-value = 0.011). In an adjusted linear model, weighted GRS was not separately associated with 3MSE scores (p-value = 0.964).

Additionally, in a linear model adjusted for the baseline demographic and clinical characteristics, we did not find an interaction between GRS and CDR to be significantly associated with lower 3MSE scores (p-value=0.478), indicating that the association of GRS with 3MSE scores does not change depending on CDR status. However, excluding the interaction term, controlling for GRS, women with large CDR still had statistically significantly lower 3MSE scores when compared to women without large CDR, yielding a predicted mean difference in 3MSE scores of 0.84 (p-value = 0.007) (Table 4).

**Table 4.** Linear Regression Model Estimates of 3MSE Scores.

| Variable* | Estimate (95% CI) | P-value |
| --- | --- | --- |
| GRS | -0.01 (-0.09, 0.07) | 0.810 |
| Large CDR | 0.15 (0.04, 0.26) | 0.007 |
| Age | 0.03 (0.02, 0.04) | <0.001 |

\*Model also adjusts for education, smoking, diabetes, BMI, cardiovascular disease, diabetic retinopathy, and hormone therapy randomization

## Discussion

This study represents the first epidemiologic investigation of the effect of GRS for vertical CDR on the relationship between optic nerve cupping and cognitive function. There are three key findings. First, a higher CDR GRS was associated with increased odds of large CDR. The odds of large CDR for a one unit increase in GRS was 2.29 (95% CI: (1.21, 4.34), p-value = 0.011). Second, there was no association between the GRS for CDR and 3MSE. Third, large optic nerve cupping was associated with poorer cognitive function, independent of the GRS.

The optic disc, specifically the CDR, is commonly assessed during ophthalmic exams to detect and monitor multiple ocular diseases, such as glaucoma. Therefore, identifying factors that affect the CDR can aid in accurate prediction of eye pathologies. Epidemiological studies have identified that high IOP and low BMI are associated with variation in CDR.^13–15^ CDR is an optic nerve head structural biomarker with a strong genetic component (heritability estimates range from 48% to 66%).^16–17^ GWAS studies have shown specific loci associated with CDR, but each locus has limited predictive power. We used a GRS based on previous work by Springelkamp et al. to identify the aggregate effect that common genetic variants have on CDR. We found an association between a higher optic nerve GRS and the risk of large CDR.

A study of Lantinx individuals also found a significant association between vertical CDR and a GRS for vertical CDR, also using SNPs from GWAS data, and controlling for age, gender, central corneal thickness, IOP, and education.^18^ They found that a weighted GRS improved the discriminatory ability for POAG, with an AUC of 0.735 (95% CI: [0.701, 0.768]).^18^ In the present study, a higher GRS was associated with a higher odds of large CDR, after adjusting for age, PC-1, and PC-2.

Previous studies also investigated GRS in relation to cognitive impairment.^19^ For instance, Wollam et al. constructed a GRS based on risk alleles and performed a neuropsychiatric evaluation to determine cognitive status. Using a logistic regression, they showed that a one unit increase in GRS was associated with a nearly 4-fold increased risk of cognitive impairment (OR = 3.824, P = .013).^19^ Our work did not show a relationship between GRS of CDR and cognitive decline. However, we utilized cognitive function at a baseline 3MSE measurement. The data collection for this study lasted several years with years of longitudinal data measuring cognitive function. While 3MSE measurements were available over time, CDR classification was only available at one time point. It would be interesting to obtain longitudinal images of CDR to investigate the relationship between the GRS and MSE measurements over time.

Our study has notable strengths and limitations. We used a large, well characterized cohort of women from WHISE and WHIMS who contributed not only optic nerve cupping data but also variables associated with optic nerve cupping and cognitive function. This study builds directly on prior work in this cohort, using the 3MSE scores, a reliable measure of global cognitive function, with very high interrater reliability (0.98), internal consistency (0.91), and test-retest reliability (0.78), when measured in older adults.^11, 20^ Although, the WHISE and WHIMS included women from a variety of racial backgrounds, the constructed GRS comprised of genetic variants validated only in White individuals and therefore the study was limited to White individuals. The inclusion of all race/ethnicity groups is of utmost importance. In this study, our analysis was limited to White participants for the two main reasons. First, the majority of the WHI participants are from White race/ethnicity (86%). In addition, the number of participants that were non-white with CDR was limited, making it challenging to perform subset analyses or to adjust for race/ethnicity. Secondly, the meta- analysis, which included the target SNPs, was primarily based on a population of European ancestry. In future studies, the inclusion of multi-racial/ethnicity is needed as results in White women are not generalizable to men and other racial groups. Future studies should validate these findings in other racial groups. Our approach to computing the GRS was based on prior work but may have been limited because we did not independently identify SNPs associated with large CDR. The SNPs previously identified were associated with a continuous measure of CDR while we used a binary CDR measure, which may have decreased the power to detect an effect of each SNP individually. Additional approaches involve the computation of a genetic risk score for cognitive function. In our future work, we plan to add detailed grading of the optic nerve and vascular analysis. Lastly, in the time since data collection, more CDR SNPs have been identified.^21^ A larger CDR SNP panel might yield different results.

## Conclusions

In conclusion, the CDR GRS and structural optic nerve biomarkers have potential to be powerful biomarkers in medicine and public health efforts. Specifically, the GRS may identify populations at risk for pathologies such as high CDR and cognitive decline. This study confirmed that a CDR GRS can predict large CDR, and the large optic nerve cupping was associated with cognitive decline, independently of the GRS for the CDR. Future research examining these biomarkers and their clinical utility with a long-term follow up is warranted.

## Acknowledgments

Dr. Hallak and Dr. Vajaranant contributed equally as co-senior authors

